# A dual functioning small RNA/Riboswitch controls the expression of the methionine biosynthesis regulator SahR in *Desulfovibrio vulgaris* Hildenborough

**DOI:** 10.1101/803072

**Authors:** M. L. Kempher, A. S. Burns, P. S. Novichkov, K. S. Bender

## Abstract

Riboswitches are *cis*-acting RNA regulatory elements that control expression of a downstream gene(s) by directly binding to a specific metabolite. Here we report a *S*-adenosylmethionine (SAM)-I riboswitch in the sulfate-reducing bacterium *Desulfovibrio vulgaris* Hildenborough (*Dv*H) that plays an additional regulatory role as a *trans* small noncoding RNA (sRNA) targeting the methionine biosynthesis cycle transcriptional regulator SahR. Sequence and expression analyses indicated that DseA (*Desulfovibrio S*AM *e*lement *A*) is located upstream of a small hypothetical protein DVU1170 and that the two are co-transcribed. Multiple techniques were used to verify the riboswitch activity of DseA and its activity as a transcriptional terminator in response to SAM. While determining a potential role for DseA in the methionine biosynthesis pathway, a mRNA target encoding SahR was identified. Subsequent electrophoretic mobility shift assays confirmed the ability of DseA to bind the *sahR* transcript and qRT-PCR analysis of a DseA deletion strain suggested a negative regulatory role. This study presents the first regulatory role for a newly discovered sRNA in *Desulfovibrio*. Additionally, this study suggests that DseA acts not only as a riboswitch, but also as a *trans* regulatory molecule.

**IMPORTANCE:** Sulfate-reducing bacteria (SRB) are important contributors to global geochemical cycles while also causing major issues for the petroleum and oil industry due to biocorrosion and souring of oil wells. Despite their significance, gene regulatory networks and pathways remain poorly understood in SRB. Here, we report a *trans* acting small noncoding RNA that plays a dual role as a SAM sensing riboswitch that controls the expression of a small hypothetical protein. Our findings provide important insights into the regulatory repertoire of sulfate-reducing bacteria.

## INTRODUCTION

Microorganisms living outside of the laboratory environment encounter an ever-changing landscape of nutrient levels and physiological conditions. Bacteria have evolved intricate systems for regulating their internal response to external stimuli to ensure survival. The importance of regulation by protein factors has been known for many decades. However, the extent to which regulation by both *trans* and *cis* acting RNA has recently become more evident as the use of RNA to regulate expression allows for a quick and transient response to environmental cues in comparison to regulatory proteins. Thus, both small noncoding RNAs (sRNAs) and riboswitches have been shown to be extremely important players in many bacterial regulatory pathways (1–8).

The central role played by sRNAs in regulatory networks has become apparent over the last decade in many types of bacteria. In *E. coli* over 100 sRNAs have been confirmed (9,10). sRNAs have been identified and characterized in many bacterial genomes including pathogens like *Yersinia pestis*, *Francisella tularensis*, and *Listeria monocytogenes* and environmentally relevant Alphaproteobacteria like *Sinorhizobium meliloti*, *Bradyrhizobium japonicum*, and *Rhodopseudomonas palustris* (11–16). sRNAs have been shown to regulate numerous stress responses in bacteria including oxidative stress, osmotic stress, carbon starvation, iron starvation, photooxidative stress, and glucose-phosphate accumulation (17–22). Most sRNAs act post-transcriptionally by binding to target mRNAs and regulating expression. Often a sRNA base-pairs to the 5′ untranslated region (UTR) of a mRNA and blocks ribosome access inhibiting translation (19). Less reported modes of regulation include activation of translation of a mRNA by inducing secondary structure changes or sequestration of a protein target (23–26).

Beyond regulation by *trans*-acting RNA molecules, several *cis*-regulatory elements are also present in many bacteria. Riboswitches are elements usually found near the 5’ end of nascent RNA molecules that can fold into two mutually exclusive secondary structures based on cellular concentrations of certain metabolites (27,28). Which structure is formed determines the fate of a downstream gene(s) by either forming a transcriptional terminator or affecting translation initiation. Many classes of riboswitches are known that sense a number of different small metabolites. The genes that are controlled by riboswitches are generally involved in the synthesis or uptake of the metabolite being sensed (27,28).

Six different classes of *S*-adenosylmethionine (SAM) sensing riboswitches have been implicated to control genes of the methionine biosynthesis cycle in different bacteria (29,30). Methionine is not only important for protein initiation and synthesis, but it is also the precursor to SAM. SAM is important in the bacterial cell as a methyl donor in a myriad of reactions by methyl transferases that are important in nucleic acid, protein, and lipid modifications. The end product of methylation reactions is s-adenosylhomocysteine (SAH), which binds to methyltransferases with greater affinity than SAM and thereby inhibits their ability to further methylate (31). It is for this reason that it is important to quickly recycle SAH and keep levels of SAM high enough to carry out necessary methylation reactions.

Many bacteria can make methionine *de novo* (31,32). This pathway in well-studied bacteria is tightly regulated through a complex feedback loop. Some bacteria do not employ any riboswitches to control methionine biosynthesis and instead use both activator and repressor proteins that have been shown to interact with SAM, SAH, or homocysteine (31–33). Despite the breadth of characterized regulators in the methionine pathway, many bacteria lack annotated riboswitches or homologs to any identified methionine regulatory proteins. Of particular importance to this study is the recently identified transcriptional regulator SahR, which seems to play a role in regulation of methionine biosynthesis genes in both Alphaproteobacteria and Deltaproteobacteria (34).

Recently, we identified several novel sRNAs in the sulfate-reducing bacterium *Desulfovibrio vulgaris* Hildenborough (*Dv*H) using high-throughput RNA sequencing. Determining the exact biological roles of these candidates has lagged behind due to limited molecular tools in *Dv*H. While investigating the potential role of one of the candidates it was determined to contain a SAM-I riboswitch domain. Here we report a novel *trans*-acting sRNA, DseA (*Desulfovibrio S*AM *e*lement *A*), which was found to not only target the mRNA of the newly discovered transcriptional regulator of SAM cycle genes (SahR), but to also function as a SAM sensing riboswitch for a small hypothetical protein. This is the first report linking a sRNA in *Dv*H to a regulatory role and a further step in the elucidation of methionine biosynthesis in *Desulfovibrio*. These findings expand our knowledge of the repertoire of regulatory mechanisms utilized by *Dv*H.

## RESULTS

### Identification of DseA sRNA

A previous study in this laboratory identified potential sRNA candidates using high-throughput RNA sequencing (manuscript in preparation). The predicted coordinates of candidate DseA were 1,264,233 – 1,264,174 on the negative strand of the *Dv*H genome. Northern blot analysis using a 30-mer probe targeting this region verified the expression of a transcript around ~150 nt in both exponential and stationary growth phases (Figure 1A). Circular RACE was performed on *Dv*H RNA with primers specific for DseA (Table S2). The majority (90%) of RACE clones sequenced determined DseA to be 164 nt in length with coordinates of 1,264,319 – 1,264,156 on the negative strand (Figure S1). The 3’ end of the determined transcript contains an inverted repeat and a string of uracils, which is common for intrinsic terminators (Figure 1B). Consensus sequences of promoters have been identified for *Dv*H for the sigma factors σ^70^, σ^54^, and σ^28^ (35). Upon visual inspection it was determined that DseA contains a promoter for σ^70^ that comprises 9% of the total promoters thus far identified in *Dv*H (Figure 1B; (35)). A small hypothetical protein (DVU1170) and a methyl-accepting chemotaxis protein (DVU1169) are located downstream of DseA, while an integral membrane protein (MviN-1) is located upstream on the opposite strand (Figure 1C; MicrobesOnline (36)).

**Figure 1.**
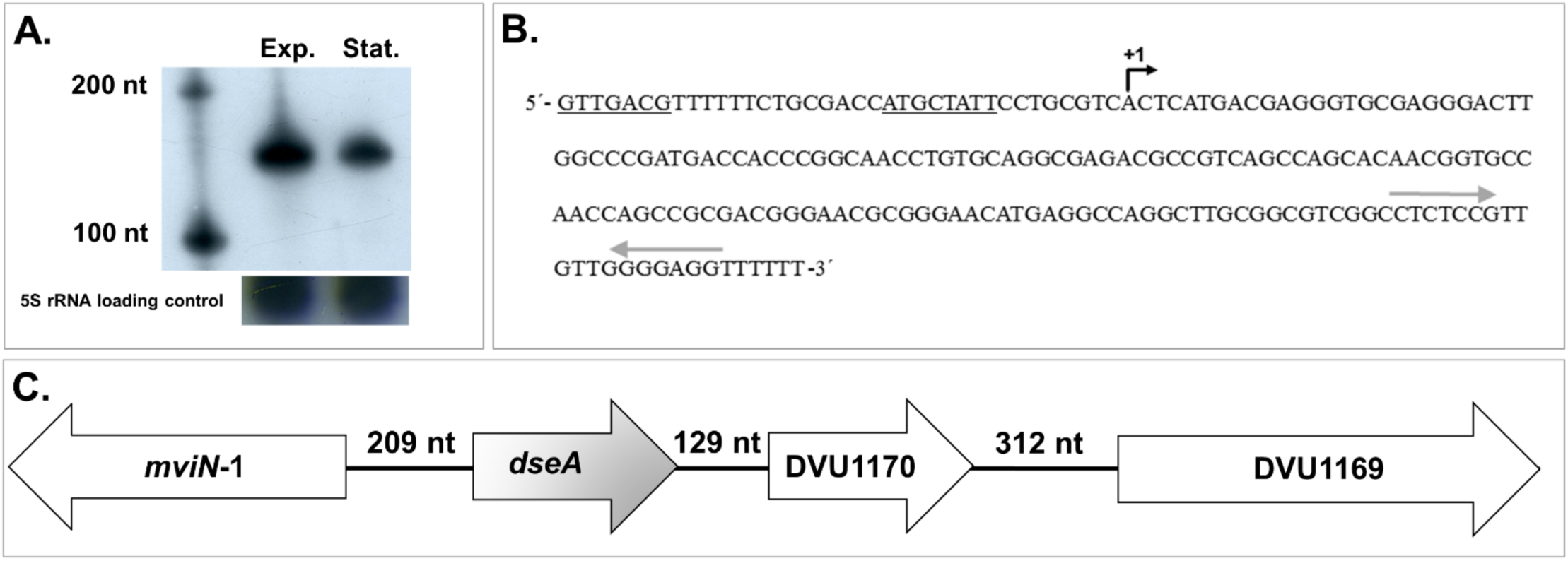
Expression of DseA and chromosomal location. (**A**) Northern Blot analysis showing 10 µg of *Dv*H RNA from both exponential (Exp.) and stationary (Stat.) growth phases. The blot was hybridized with the DseA 30mer probe (Table S2) and size was determined by comparison to RiboRuler™ Low Range RNA ladder. The membrane was stripped and re-probed for the 5S rRNA. (**B**) The DNA sequence of DseA determined by circular RACE. The predicted σ^70^ promoter is underlined. The +1 site of transcription is indicated with a black arrow. The inverted repeats of the predicted intrinsic terminator are indicated with grey arrows. (**C**) Genome view of the region around DseA encompassing coordinates 1,266,111 – 1,263,449. Gene size and location are not to scale.

### Conservation of DseA and Identification of a SAM-I riboswitch domain

The NCBI Basic Local Alignment Search Tool (BLAST) was used to search for conservation of DseA in other organisms. Similar sequences were found in *Desulfovibrio alaskensis* G20 (82% identical), *Desulfovibrio salexigens* DSM 2638 (83% identical), and *D. vulgaris* strains RCH1 (100% identical), DP4 (100% identical), and Miyazaki F (76% identical) (Figure S2). Noncoding RNA elements can often have similar structures regardless of primary structure. Therefore, the Rfam database (37) was used to search for structural homologs of DseA. This search resulted in the identification of a SAM-I riboswitch element within DseA (Figure S3). This predicted SAM riboswitch element contained an intrinsic terminator and suggested the mode of regulation to be that of transcription termination. Therefore, a small transcript would result during the “off” state and a longer transcript would result from read-through of the terminator in the “on” state.

Because a small transcript had already been confirmed by Northern blot analysis, but a larger transcript had not been observed using our methods, we further investigated the potential of DseA to act as a SAM-I riboswitch controlling the expression of the downstream gene DVU1170. Reverse-transcriptase PCR (RT-PCR) was used to confirm the presence of a larger transcript corresponding to read-through of the predicted DseA terminator into the 189 nt DVU1170 (located 129 nt downstream of the predicted riboswitch; Figure 1C). Additional primer sets spanning the region between DseA and DVU1170 confirmed co-transcription (Figure S4A). Additionally, DVU1170 was not predicted to be part of an operon and this was confirmed by RT-PCR using primers designed to anneal to the downstream gene DVU1169 (Figure S4B).

### Expression of DVU1170

The read-through transcript of DVU1170 was impossible to visualize via Northern Blot analysis. Therefore, qRT-PCR was used to monitor the presence of the read-through transcript of DVU1170 during growth with or without added methionine. SAM is synthesized from methionine and an increase in methionine levels has been shown in other bacteria to lead to an increase of SAM levels inside of the cell (38). Additionally, it is unknown if *Dv*H can uptake SAM directly from the growth medium despite the computational prediction of a methionine transporter (31). Therefore, qRT-PCR was performed on RNA extracted from *Dv*H cultures that had been grown to early exponential phase, separated into two, and then either spiked with 1 mM methionine or an equal volume of degassed H_2_O. RNA was taken at 5, 15, 30, 60, and 120 min post separation. qRT-PCR was done using RNA from each time-point and normalized using the 16S rRNA transcript. Levels of DVU1170 transcript dropped after the addition of methionine by 2.97-fold at 5 min and up to a 3.47-fold decrease at 15 min compared to the sample without methionine at the same time-point (Figure 2). These data suggest that levels of DVU1170 expression are influenced by concentrations of methionine.

**Figure 2.**
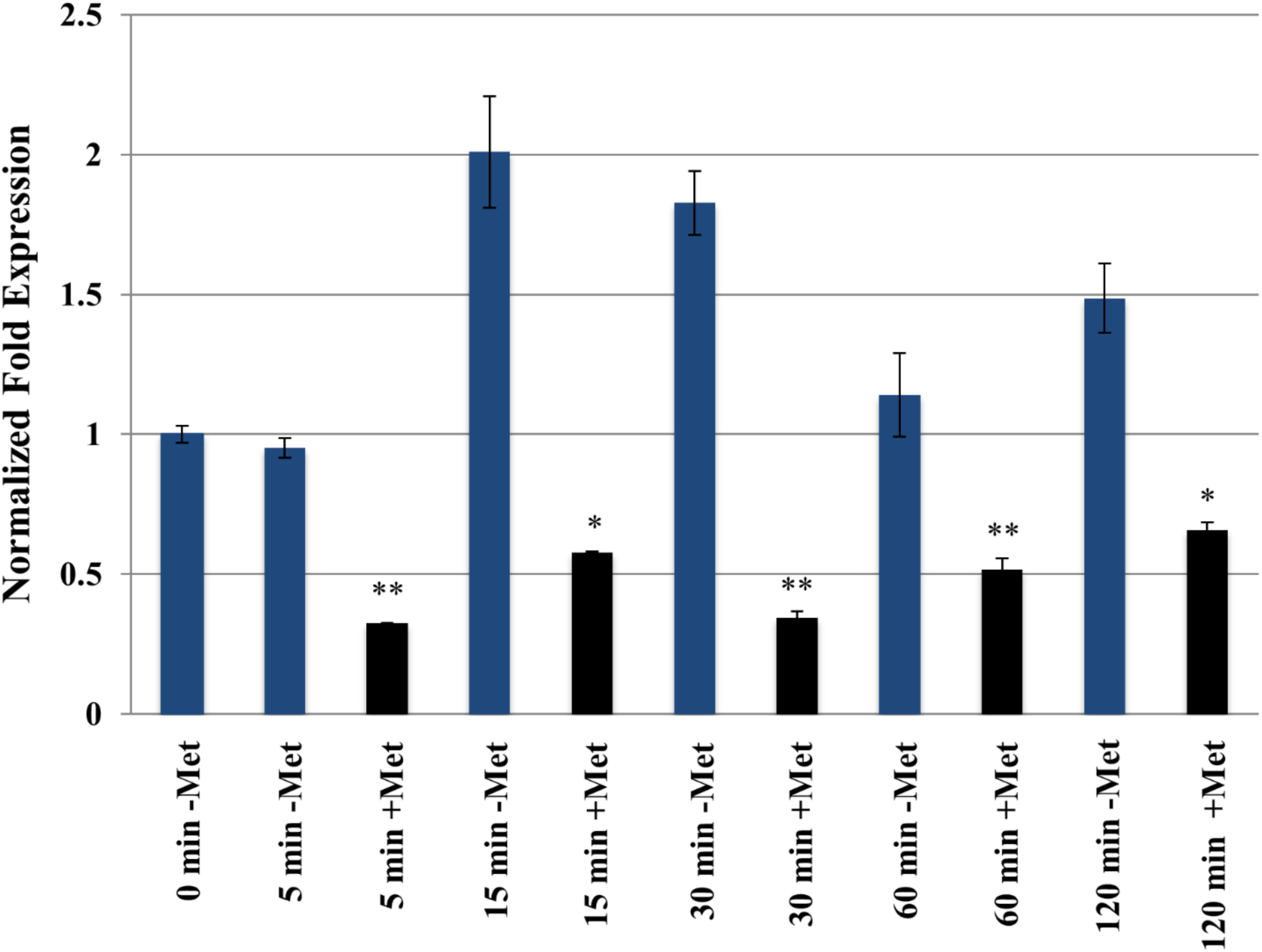
Relative quantification of DVU1170 during growth with or without added methionine. Transcript levels were normalized to the 16S rRNA transcript (using primers DVU1170 qRT-PCR F/R and 16S qRT-PCR F2/R2; Table S2). Expression of the 0 min control was artificially set to 1 and expression data for the remaining time-points were determined by the CFX™ Manager software. Error bars represent standard error. Samples with added methionine were compared to the same time-point without methionine using Student t test, two tailed (* p< 0.05, ** p< 0.01).

### Direct binding of SAM to the DseA riboswitch

SAM riboswitches directly bind SAM while discriminating against very structurally similar compounds like methionine and SAH. Upon binding by SAM, structural rearrangements occur in the expression platform (39). The first 165 nt of the DVU1170 UTR was *in vitro* transcribed and subjected to in-line probing which reveals locations of structured versus unstructured RNA based on differing rates of spontaneous cleavage of RNA. RNA phosphodiester linkages are cleaved by the ribose 2’ oxygen on the adjacent phosphorus and the rate of this reaction depends on the “in-line” position of the 2’ oxygen, phosphorus, and 5’ leaving group (40). RNA samples were mixed with 0.1 mM, 0.5 mM, and 1 mM SAM and compared to a sample with no additional factors, 1 mM methionine, or 1 mM SAH. There were several regions of the RNA where scission either increased or decreased based on the addition of SAM indicating a difference in structural conformation (Figure 3A). The areas indicated with an arrow labeled 1, 2, 4, or 5 had increased scission in the presence of SAM based on lane profiles obtained for each sample while arrows marked with 3, 6, or 7 indicated decreased scission in the presence of SAM. No differences were observed between the negative control, the methionine sample, or the SAH sample. The lane profiles were normalized based on total signal and then plotted as intensity versus lane position (Figure 3B). These findings validate that the structural rearrangement is specific to SAM and not affected by methionine or SAH.

**Figure 3.**
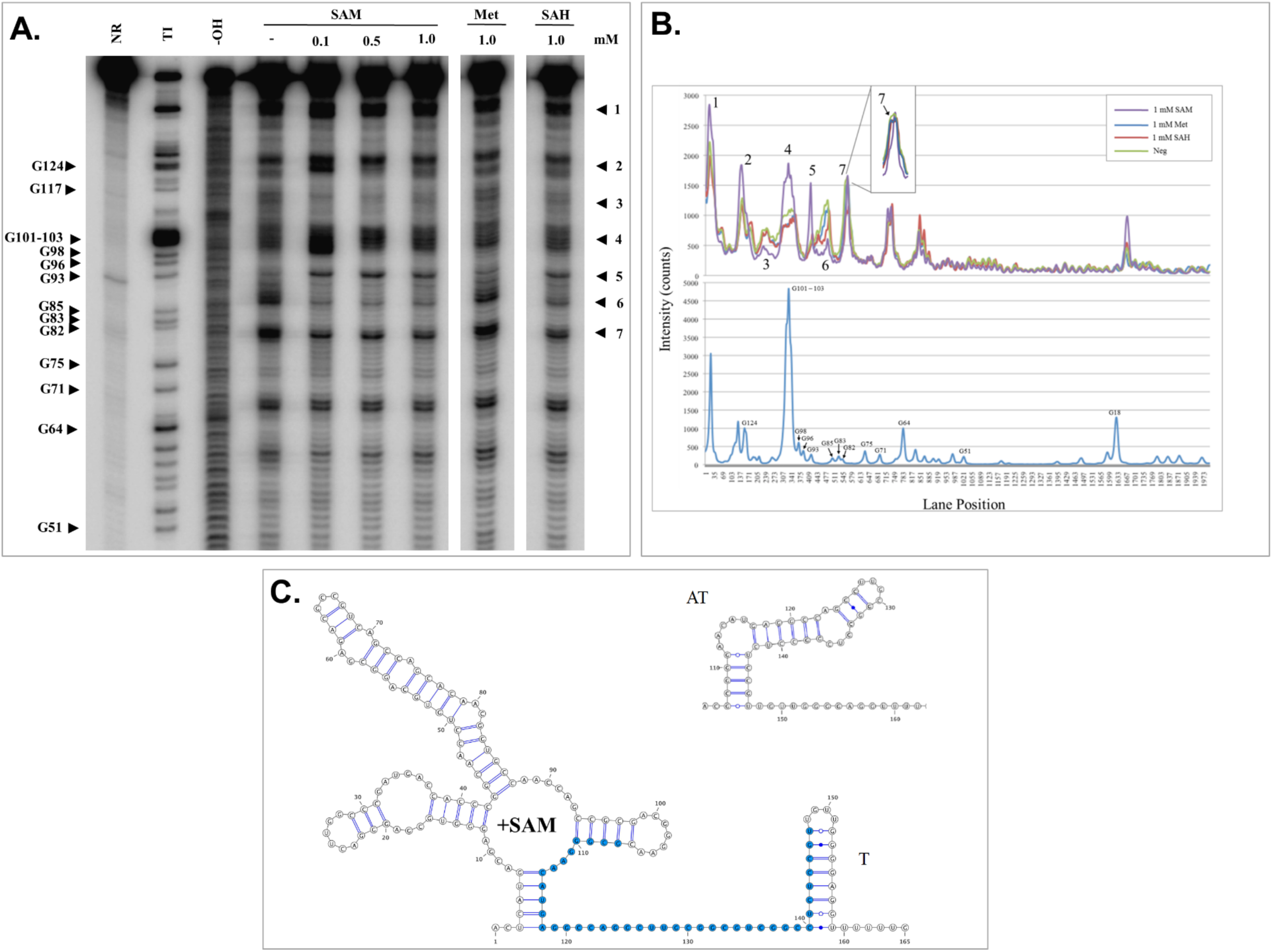
Structural analysis of the DseA riboswitch. (**A**) Spontaneous cleavage pattern of DseA in the absence or presence of SAM, methionine (Met), or SAH as indicated. The location of some of the guanosine residues (G) cleaved by RNase T1 is indicated. NR: no reaction; T1: RNase T1 ladder; -OH: alkaline hydrolysis ladder. (**B**) Lane profiles as determined by the program ImageQuant (GE Healthcare) of in-line probing gel. The numbers match to the same numbered areas of the gel. The lane profile of the T1 ladder is plotted in the bottom panel and represents the G residues as labeled. (**C**) Predicted secondary structure of DseA. When SAM concentrations are high an intrinsic terminator (T) is predicted to form. When SAM concentrations drop the anti-terminator (AT) forms instead. Bases colored blue are involved in forming the antiterminator.

The data from the in-line probing analysis were used to draw a probable secondary structure using the RNA structure visualization program VARNA (41). The structural model is consistent with other characterized SAM-I structures and indicated an anti-terminator to form in the absence of SAM and a terminator stem loop to form in the presence of SAM (Figure 3C; (42)).

### SAM promotes transcription termination *in vitro*

Previous studies have indicated that riboswitches can act at the level of transcription termination or translation initiation (27,43). While the predicted DseA riboswitch region contains a characteristic intrinsic terminator (G + C rich region followed by a series of uracils), suggesting that the riboswitch functions at the level of transcription termination, single-round *in vitro* transcription termination assays were carried out to determine if the addition of SAM affected the amount of transcription termination. These assays were performed by linking the *Dv*H DseA region, including the predicted terminator and ~184 nt downstream, to a T7A1 promoter recognized by *E. coli* RNAP. If termination does not occur, then the RNA polymerase will continue to transcribe the DNA until it reaches the end of the template and falls off (“read-through” transcription).

A single mixture of halted complexes was created before being separated and mixed with various amounts of SAM or methionine. When no additional factors were added to the mixture a termination rate of 52% was observed (Figure 4A, lane 1). The termination frequency increased to 63% and 82% when 0.1 mM and 0.5 mM of SAM was added, respectively (Figure 4A, lanes 2 and 3). When 0.5 mM of methionine was added a termination rate of 55% was observed which was similar to the negative control (Figure 4A, lane 4). These findings indicate that upon SAM binding, the anti-terminator sequence is sequestered, and an intrinsic terminator is formed instead halting transcription. These data also indicate the response to SAM is specific since methionine had no effect on termination.

**Figure 4.**
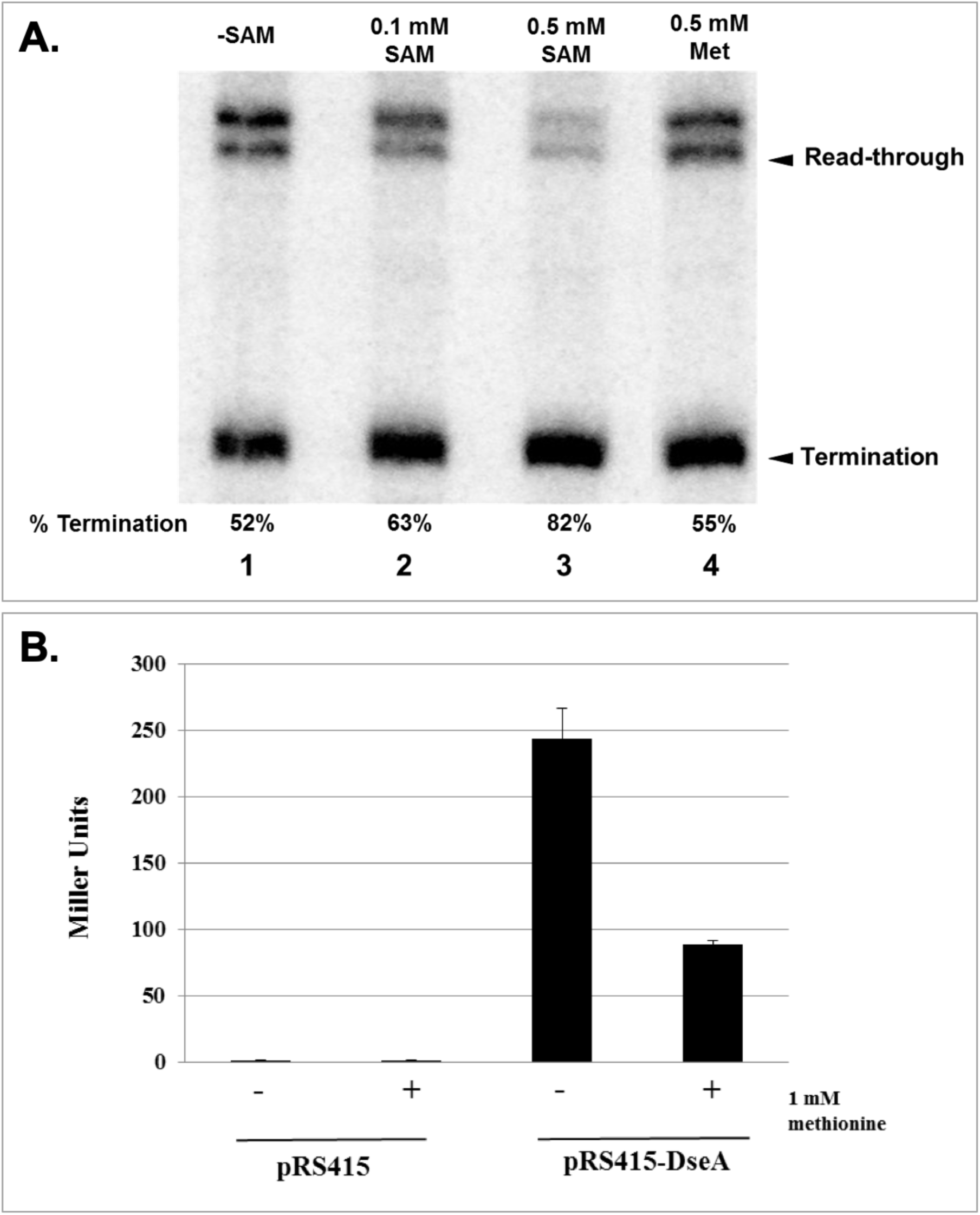
DseA riboswtich expression platform response. (**A**) *In vitro* transcription termination assay of the riboswitch region. Percent termination was determined by the amount of termination product divided by the sum of total transcription products. (**B**) β-galactosidase activity of pRS415-DseA and the negative control vector pRS415. Values represent the mean of three experiments. Activity is represented by Miller Units. Error bars represent standard deviation.

### Transcriptional fusion of the DseA promoter and predicted riboswitch to *lacZ*

SAM riboswitches have shown a large range of variation in their response to different levels of SAM *in vitro* (38). Transcriptional *lacZ* fusions were made to determine if transcription termination also increased *in vivo*. Currently, no reporter gene system is available for *Dv*H. Therefore, the fusions were made in an *E. coli* background. The predicted promoter for DseA and the entire riboswitch region were ligated into pRS415 directly in front of a promoterless *lacZ* gene and transformed into *E. coli*. As *E. coli* cells are unable to uptake SAM directly, methionine was used to act as a SAM precursor. The strain containing the DseA-*lacZ* fusion and a control strain with empty vector were grown in minimal media containing 1 mM methionine to early exponential phase, separated into two different cultures, centrifuged, and resuspended in minimal media with or without 1 mM methionine. After 3 h of growth the samples were assayed for β-galactosidase production.

The negative control strain containing the empty pRS415 vector produced 1.09 Miller Units (MU) without added methionine and 1.32 MU with added methionine. The strain containing the pRS415-DseA vector produced 230.48 MU without additional methionine and 88.7 MU with added methionine indicating a 2.5-fold decrease in β-galactosidase production after the addition of methionine (Figure 4B). These findings provided further evidence that transcription of DVU1170 is regulated in response to methionine concentrations in *E. coli*.

### Prediction of a mRNA target of the DseA sRNA

In all the conditions tested (rich media, minimal media, H_2_O_2_ stress, salt stress, and cells grown in a biofilm) the terminated DseA product was abundant, even after several hours of growth (data not shown). This indicated that perhaps the terminated product was playing an additional role. One riboswitch has been shown to also act in *trans* as a sRNA inhibiting a mRNA target (44). Thus we used the computational prediction program IntaRNA to compile a list of potential mRNA targets of DseA (Table S3; (45)). One of the top target candidates was the mRNA for SahR (DVU0606), which had recently been shown to act as a transcription factor that controls the known methionine biosynthesis and SAM cycle genes in *Dv*H and other Deltaproteobacteria (34).

Since SAM riboswitches are known to play a role in the methionine biosynthesis cycle in other bacteria we decided to investigate the possibility of DseA targeting the *sahR* mRNA. The predicted base pairing between DseA and the *sahR* mRNA blocks the ribosome binding site and the start codon likely inhibiting translation of *sahR* (Figure 5A).

**Figure 5.**
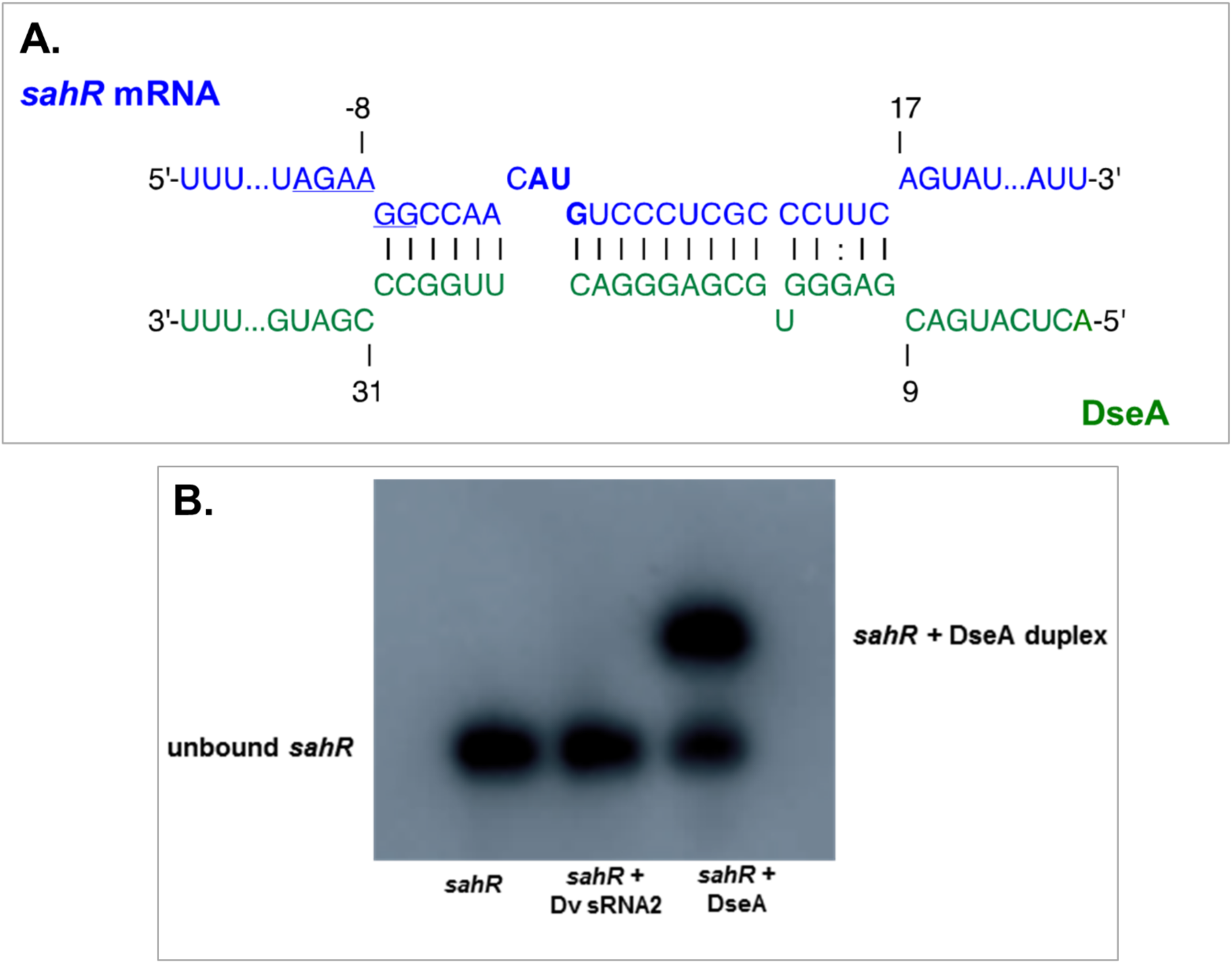
Interaction of DseA with s*ahR* RNA. (**A**) Predicted interaction region of DseA (green) and *sahR* (blue) as determined by IntaRNA (55). The RBS is underlined while the start codon is bolded. (**B**) EMSA showing radiolabeled *sahR* alone, mixed with Dv sRNA2, and DseA. The migration of free *sahR* and bound *sahR* is indicated. The following primers were used to generate *in vitro* transcripts: SahR T7 prom F/R, Dv sRNA-2 T7 prom F/R, and DseA T7 prom F/R (Table S2).

### DseA sRNA and *sahR* 5’ UTR directly interact *in vitro*

Full length DseA and the 5’ UTR of *sahR* were synthesized to validate a direct interaction between the two molecules using an electrophoretic mobility shift assay (EMSA). The 5’ portion of *sahR*, corresponding to the −31 to +168 nucleotides relative to the start codon of *sahR* mRNA, was *in vitro* transcribed and radiolabeled at the 5’ end. DseA was also *in vitro* transcribed but was not radiolabeled. The two RNA molecules were mixed together and then analyzed by native-PAGE analysis. A sample with only the *sahR* RNA showed the migration pattern of free *sahR*. When DseA was added this band shifted up confirming the interaction of the two RNA molecules (Figure 5B). Another sRNA identified in a previous study was used as a control RNA (Dv sRNA2) and no shift was seen when this RNA was added with *sahR* instead of DseA.

An additional EMSA analysis was done on smaller portions of the *sahR* mRNA to determine the exact nucleotides involved in binding (Figure S5). An RNA oligo that corresponded to the nucleotide positions of −31 to +46 relative to the start codon of *sahR* mRNA was sufficient to bind to DseA and show a shift. A region that corresponded to the nucleotide positions +17 to +88 relative to the start codon of *sahR* mRNA did not shift. This confirmed that the predicted region of interaction near the RBS and start codon of *sahR* is required for interaction with DseA.

### DseA sRNA controls expression of *sahR* mRNA

Since DseA was predicted to bind to the RBS of the *sahR* mRNA, it seemed likely that the mode of regulation of the sRNA would be to block translation of *sahR* which can often lead to degradation of the targeted mRNA (46,47). To investigate this, the levels of *sahR* transcript were evaluated in a DseA deletion mutant (Δ*dseA*) and compared to wild-type levels. qRT-PCR was done on RNA extracted from both exponential growth and stationary growth of wild-type *Dv*H and the Δ*dseA* strain. The level of the *sahR* transcript was higher in the Δ*dseA* strain suggesting that the mode of action of DseA is inhibitory (Figure 6A). The expression of *sahR* increased 1.48-fold during exponential growth and almost 20-fold during stationary growth in the Δ*dseA* strain. In order to exclude the possibility that the changes in expression were due to inactivation of the downstream gene DVU1170, a complement strain was constructed that expressed only DseA from a constitutive promoter. Expression from this promoter resulted in higher levels of expression of DseA than compared to the wild-type strain. Expression of *sahR* was significantly down regulated in the complement strain further suggesting an inhibitory role for DseA.

**Figure 6.**
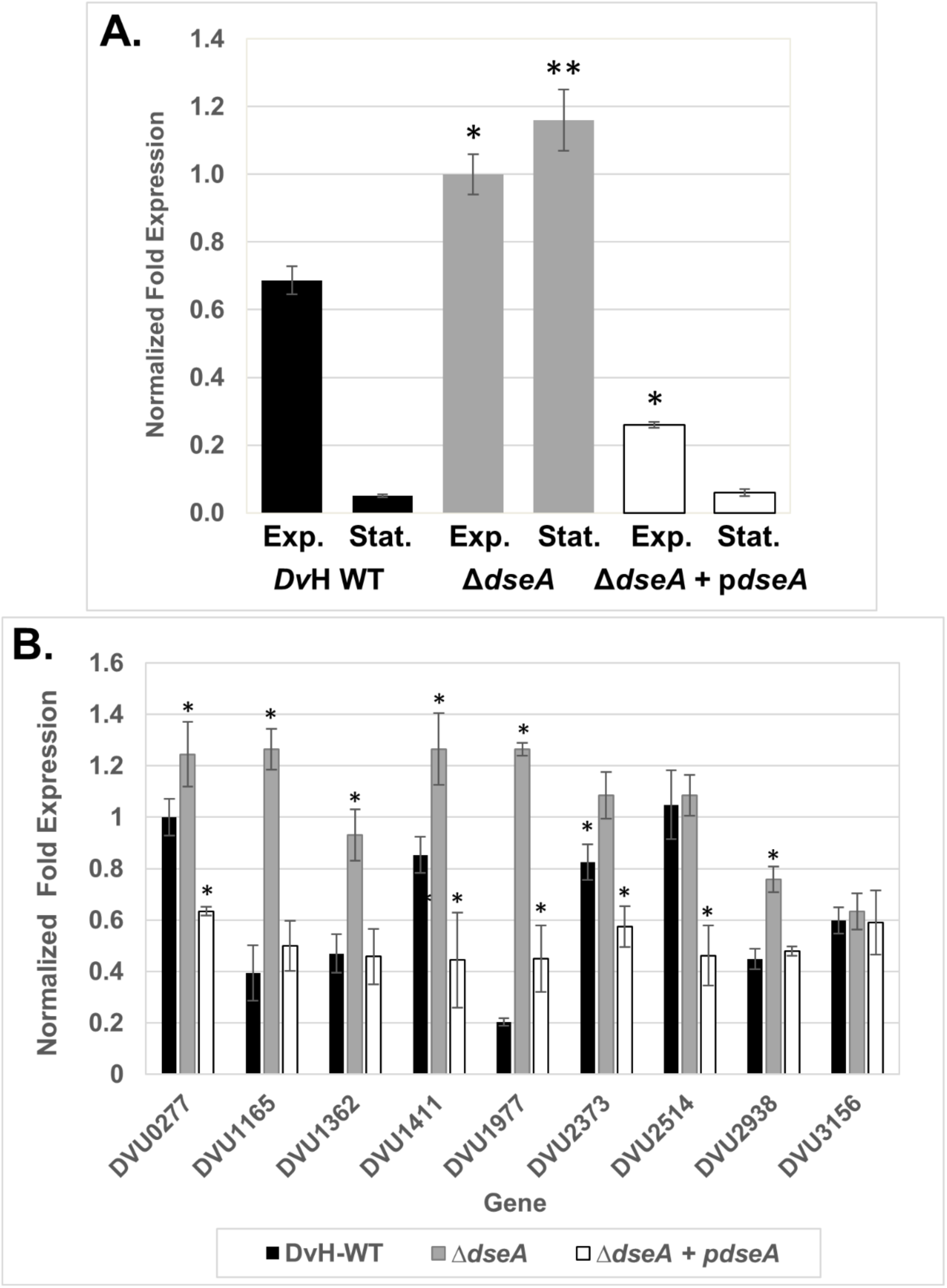
**(A)** qRT-PCR analysis of *sahR* expression in both exponential (Exp.) and stationary (Stat.) growth. **(B)** qRT-PCR analysis of additional predicted targets of DseA during exponential growth. Transcript levels were determined for wild-type DvH, **Δ***dseA*, and the complement (**Δ***dseA* + p*dseA*). Each gene was normalized to the 16S rRNA and *rplS* reference genes. The efficiency of each primer pair is as follows: *sahR*- 90.1%, DVU0277-88.9%, DVU1165- 90.0%, DVU1362-89.7%, DVU1411-89.5%, DVU1977- 85.2%, DVU2373-90.2%, DVU2514-91.3%, DVU2938-94.9%, DVU3156-89.8%, 16S rRNA gene-92.5%, *rplS*-97.0%. Error bars represent standard error. Samples from **Δ***dseA* and **Δ***dseA* + p*dseA* were compared to the wild-type sample from the same growth phase using Student t test, two tailed (* p< 0.05, ** p< 0.01).

### Additional putative targets of DseA

Typically, sRNA-target interactions are confirmed *in vivo* via systems that link expression of the mRNA target to a reporter gene (either chromosomally or on a plasmid) and then placing the sRNA behind an inducible promoter on a vector (48). Unfortunately, no such system exists for *Dv*H. We, therefore, attempted to heterologously express DseA and *sahR* using a system in *E. coli* where the mRNA target is chromosomally inserted behind an inducible promoter and translationally fused to *lacZ* in strain PM1205 (49,50). The sRNA is then expressed behind an inducible promoter from a vector. However, we were unable to get appreciable levels of expression of the *sahR-lacZ* fusion (~12% of the amount we were able to obtain with the 5’ end of the *E. coli manX* mRNA fused to *lacZ*; data not shown), possibly due to slight differences in ribosome binding sites between *E. coli* and *Dv*H. Therefore, to provide further evidence to support the putative role of DseA as a regulatory sRNA, we performed qRT-PCR on additional IntaRNA predicted mRNA targets (Table S3). Eight of the top predicted targets were selected for analysis. Additionally, the entire list of predicted targets was manually searched for any genes that had been identified as part of the methionine biosynthesis pathway. A recent study identified that a DUF39 protein was required for homocysteine formation in *Desulfovibrio alaskensis* (51). The homolog of this gene in *Dv*H, DVU2398, was a predicted target of DseA and was included in the qRT-PCR analysis. Of the nine genes analyzed, seven showed a significant difference in expression between the wild-type and the *ΔdseA* strain (Figure 6B). Furthermore, expression of the affected genes was either restored to similar levels observed in the wild-type strain or down-regulated to a greater extent in the complement strain indicating the observed effect was due to DseA and not the downstream gene DVU1170.

## DISCUSSION

Regulation of gene expression by both *trans* acting sRNAs and *cis* acting riboswitch elements has been implicated in numerous nutritional regulatory networks. This study is the first in *Desulfovibrio* to both examine the mechanism of a SAM riboswitch and suggest a definitive role for a sRNA.

While a previous study had predicted the presence of riboswitches such as thiamine and vitamin B_12_ elements in *Desulfovibrio*, no SAM riboswitch was identified (33). The DseA element was likely overlooked in this study as only regions linked to genes predicted to be involved in methionine biosynthesis were analyzed. Characterized SAM-I riboswitches from other bacteria have been shown to bind SAM but discriminate against methionine and other similar metabolites (39). Results from the in-line probing assay suggested changes in secondary structure occur when SAM is present but not when methionine or SAH is present (Figure 3). The specificity of the RNA transcript to bind to SAM but not to methionine or SAH agrees with previous evidence that SAM riboswitches are highly specific to SAM as their sole metabolite. Analysis of *in vitro* transcription termination in the presence or absence of SAM assays also indicated that a significant increase in termination occurred when SAM was added but not in the presence of methionine (Figure 4A). This provides further evidence that the DseA riboswitch is specific for SAM and that it acts at the level of transcription termination.

Transcriptional *lacZ* fusions in *E. coli* further corroborated that *in vivo* changes occurred in response to methionine concentrations as samples without methionine showed greater β-galactosidase activity compared to samples with methionine (Figure 4B). These data suggest that increased levels of methionine lead to lower levels of the downstream gene. Even with added methionine the complete inhibition of β-galactosidase was not seen. Perhaps tighter control would be seen with greater amounts of methionine or with *in vivo* studies in *Dv*H as opposed to *E. coli*.

In this study we showed that DseA can also bind to *sahR* (DVU0606) mRNA *in vitro*, providing further support that DseA acts as both a SAM responsive riboswitch and a *trans* acting sRNA (Figure 5). The region surrounding the RBS and start codon of the *sahR* transcript is necessary for DseA binding to occur and this suggested the mode of regulation to be inhibitory (Figure S5). Comparison of *sahR* transcript levels in a DseA deletion mutant to those of the wild-type supported this hypothesis (Figure 6). RNA extracted from the deletion strain of DseA showed an almost 20-fold increase in the level of expression of *sahR* during stationary phase. Seven additional predicted targets of DseA showed a similar pattern of increased expression in the DseA deletion strain. Of particular interest was the increased expression of DVU2938 in the DseA deletion strain. DVU2938 homologs in two separate *Desulfovibrio* species have recently been shown to be involved in the methionine biosynthesis pathway (51,52). These recent studies suggest that the corresponding proteins in *D. alaskensis* G20 (Dde_3007) and *D. miyazaki* (DVMF_1464) can transfer a sulfur group to *O*-phosphohomoserine to form homocysteine in the pathway for methionine biosynthesis. Thus, providing further evidence for a regulatory role of DseA related to the methionine biosynthesis pathway.

While it remains to be seen whether the change in expression observed for predicted DseA targets is due to direct binding of DseA or from a downstream effect from other regulators controlled by DseA, it is clear that DseA is playing some role in altering the expression of these genes. Whether or not a link between these additional predicted targets and regulation of the methionine biosynthesis cycle exists is beyond the scope of this present study. However, we do aim to explore the global regulatory role of DseA in future studies.

It should be noted that in the *Desulfovibrio* species in which DseA is conserved, the riboswitch is linked to homologs of the hypothetical protein DVU1170 (Figure 1C). We confirmed that DVU1170 is co-transcribed with DseA in *Dv*H and that transcript levels of DUV1170 are higher in the absence of exogenous methionine (Figures 2 and S4A). This longer transcript was only slightly visible in Northern blot analysis compared to terminated DseA under every growth condition tested. It remains to be seen if DVU1170 expression is always low compared to DseA or if the condition in which expression increases was not established in this study. While it is tempting to predict a novel role for DVU1170 in the methionine biosynthesis pathway of *Desulfovibrio*, a more focused study targeting the activity of DVU1170 will need to be done before a role for the protein *in vivo* can be determined.

Based on this study and previous data showing that SahR negatively regulates genes in the SAM cycle (34), we have constructed the model presented in Figure 7. This model suggests that when SAM concentrations are high, more premature transcription termination occurs before reaching the downstream DVU1170 gene. This would increase the levels of DseA as a *trans* acting sRNA, allowing DseA to bind to *sahR* mRNA and alter its expression. When the cell experiences high levels of SAM, it is an indicator that very high levels of SAH will soon follow. SAH is toxic and must be eliminated quickly. Decreasing the amount of the transcription factor SahR leads to derepression of genes (*ahcY*, *metE*, and *metK*) that encode products essential for recycling SAH back to SAM. However, additional experiments will need to be carried out to verify the relationship of DseA and *sahR*.

**Figure 7.**
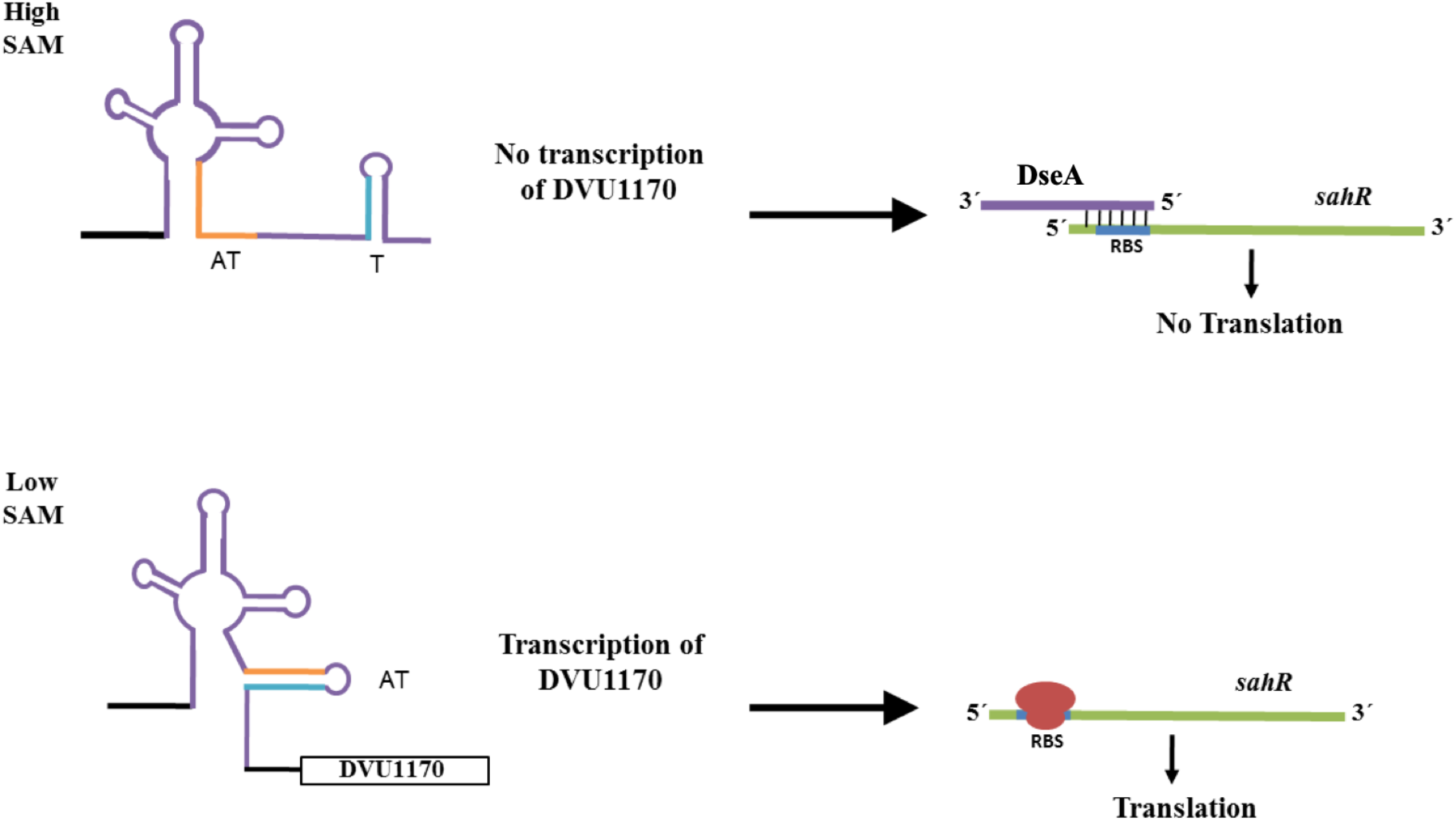
Predicted model for the activity of DseA under high SAM concentrations versus low SAM concentrations. DseA is represented in purple, the anti-terminator is orange, and the sequence shared by the terminator and anti-terminator is blue. *sahR* mRNA is shown in green. T: terminator; AT: antiterminator.

Overall, new studies are showing a myriad of regulatory roles for riboswitches (5,53). In fact, recent reports have shown that riboswitches can be used to control the downstream expression of non-coding RNAs. In *L. monocytogenes*, a vitamin B_12_ riboswitch controls the expression of an antisense RNA that targets the mRNA of the PocR transcriptional regulator (54), while in both *Enterococcus faecalis* and *L. monocytogenes* a vitamin B_12_ riboswitch also controls the transcription of trans-acting sRNAs EutX and Rli55, respectively (55,56). Full-length EutX and Rli55 possess structures that bind antiterminator proteins. When vitamin B_12_ is present, transcriptional termination occurs preventing the synthesis of full-length EutX and Rli55. These truncated sRNAs are unable to sequester antiterminator proteins and thus transcriptional read-through of ethanolamine utilization (*eut*) genes (whose products require vitamin B_12_ as a cofactor) is allowed. Our work adds to the unique and diverse repertoire of riboswitches and the multiple layers of control bacteria employ to regulate basic metabolic pathways as to our knowledge, this is only the second report of a riboswitch that plays a dual role in *trans* by inhibiting translation of a mRNA target (44). Interestingly, the other dual-acting riboswitch/sRNA is also a SAM-I riboswitch that is upstream of an ABC transporter operon in *Listeria monocytogenes*. In *trans*, the SreA sRNA decreases the level of the PrfA virulence regulator when SAM is present by negatively regulating the translation of the *prfA* mRNA (44).

It is likely that other regulators are involved in *Desulfovibrio* methionine biosynthesis and the relationship between DseA and *sahR* is much more intricate than was evaluated in this initial study. It will be necessary to investigate the regulation of the methionine biosynthesis genes *in vivo* before this pathway will be fully understood. Additionally, identification of the genes responsible for completing the methionine biosynthesis cycle and determining a role for DVU1170 will add insight into the regulation of this complex biosynthesis pathway.

## MATERIALS AND METHODS

### Bacterial strains and growth conditions

Bacterial strains and plasmids used in this study are listed in Table S1. *Desulfovibrio vulgaris* Hildenborough (*Dv*H) and strains constructed from *Dv*H were grown statically at 34°C in an anoxic chamber (Coy) with an atmosphere composed of 5% H_2_/95% N_2_ in defined lactate/sulfate medium (LS4D) reduced with 5 ml per liter of an anaerobic titanium citrate solution (57). *E. coli* strains were grown at 37°C with shaking (200 rpm) in LB medium or M9 salts minimal media (58). When necessary, media was supplemented with the appropriate antibiotics at the following concentrations: ampicillin (50 μg/ml), kanamycin (50 μg/ml), geneticin (G418; 400 μg/ml), or spectinomycin (100 µg/ml).

### Nucleic Acid Isolation

Genomic DNA (gDNA) was extracted from pure cultures grown overnight using the Wizard^®^ Genomic DNA purification kit (Promega) following the manufacturer’s protocol for Gram-negative bacteria.

RNA was extracted from *Dv*H cultures grown to either exponential (OD_600 nm_ 0.30 – 0.50) or stationary growth phase (OD_600 nm_ 0.80 – 0.90). Cultures were placed on ice and ice-cold stop solution (95% ethanol/5% phenol) was added at a final concentration of 20% (v/v). Total RNA was isolated using TRI Reagent^®^ Solution (Ambion) following the manufacturer’s guidelines. The concentration and purity of the RNA was calculated by the ND-1000 NanoDrop Spectrophotometer (Thermo Scientific). RNA samples were treated with DNase using the TURBO DNA-free kit (Ambion) following the manufacturer’s protocol.

RNA transcripts were analyzed by Northern blot analysis. Each sample contained 10 µg of DNase treated RNA mixed with an equal volume of Gel Loading Buffer II (Ambion). Samples were loaded onto a precast Novex^®^ 6% or 10% TBE-urea gel and run in a X-cell Surelock™ gel rig with 1X TBE. RiboRuler™ Low Range RNA ladder (Thermo Scientific) was prepared and labeled according to manufacturer’s guidelines using [gamma- ^32^P]-ATP (6,000 Ci/mmol) (Perkin Elmer) and T4 Polynucleotide Kinase (PNK; NEB). The gel was separated from the cassette and equilibrated in 0.5X TBE buffer along with filter pads and a Nylon Charged Membrane (GE Healthcare). Electroblotting was carried out using a Trans-Blot^®^ SD Semi-Dry Transfer Cell (Bio-Rad) for 2 h at a constant current of 200 mA. The membrane was rinsed and UV-crosslinked.

The membrane was hybridized overnight at 50°C with DNA oligo probes labeled with [gamma- ^32^P]-ATP (6000 Ci/mmol) (Perkin Elmer). 20 pmol of the DNA oligo was mixed with 7 μl of [gamma- ^32^P]-ATP, 2 μl of 10X PNK Reaction buffer, 1 μl of T4 PNK (NEB), and H_2_O up to 20 μl. Oligo mixtures were incubated at 37°C for 1 h. The probes were heated to 95°C for 5 min before being added. Membranes were washed twice with 2X SSC/0.1% SDS for 5 min followed by two washes for 15 min with 0.1X SSC/0.1% SDS. The membranes were exposed to Fuji X-Ray film overnight at −80°C. The film was manually developed using Kodak developer and fixer solutions. Membranes were stripped with a 0.5% SDS solution while rotating for 1 h at 60°C. The membranes were rinsed with DEPC-H_2_O and hybridized as described above. The membranes were then probed with a DNA oligo targeting the 5S rRNA gene as a loading control. The probe was labeled as described above. Probes used for Northern blots can be found in Table S2.

### Rapid Amplification of cDNA Ends (RACE)

The start and stop sites of transcription of DseA were determined by circular RACE as described elsewhere (59,60). Briefly, DNase-treated RNA was treated with Tobacco Acid Pyrophosphate (TAP) (Epicentre) at 37°C. The 5′ monophosphate and 3ʹ free –OH ends of the RNA were ligated with T4 RNA ligase (Invitrogen) to make circular molecules. The circular transcripts were reverse transcribed using a DseA specific primer (DseA CR R; Table S2) into first strand cDNA. PCR was used on the cDNA with primers on either side of the 5′/3′ bridge region (DseA CR F and DseA CR R; Table S2). The PCR product was cloned into pJET1.2 (Thermo Scientific) and the plasmid was sequenced to determine the transcript ends.

### RT-PCR and qRT-PCR

Reverse transcriptase (RT)-PCR was carried out using ImProm-II reverse transcriptase (Promega) following the manufacturer’s protocol with 1 µg of RNA. 5 µl of the cDNA reaction mixture was used as a template in a 50 µl PCR amplification reaction mixture with corresponding forward and reverse primers (Table S2) and GoTaq DNA polymerase (Promega), as described by the supplier. For control reactions, RNA without reverse transcriptase or chromosomal DNA was used as a template.

DNase-treated RNA extracted from various conditions was used for qRT-PCR analysis. The SuperScript^®^ III First-strand Synthesis for qRT-PCR kit (Invitrogen) was used with 1 µg of DNase-treated RNA to make cDNA. 1 µl of the cDNA was used as template and reactions were carried out using the SYBR^®^ Green SuperMix (Quanta Biosciences) and a MJ MiniOpticon™ thermocycler running CFX™ Manager software (Bio-Rad). The cycling parameters consisted of an initial denaturation step of 3 min at 95°C followed by 40 cycles of denaturation at 95°C for 30 s and annealing/extension at 63°C for 30 s. After each cycle, fluorescence was recorded. A melt curve was performed at the end of each experiment starting at 63°C and concluding at 95°C (0.5°C/5 s). A no RT control indicated no DNA contamination was present. Transcript levels were normalized to the 16S rRNA gene or the *rplS* gene (DVU0835; primer sequences obtained from Christensen *et al.* (61)). and fold changes were calculated using the Pfaffl method (62). To calculate reaction efficiency of each gene-specific primer set, a standard curve using a series of diluted cDNA (6 logs of serially diluted 100 ng/μl cDNA) was generated.

### *In vitro* transcription

RNA was *in vitro* transcribed using the MEGAshortscript™ kit (Ambion) following the manufacturer’s guidelines. Standard PCR was used to generate the DNA template from *Dv*H gDNA. RNA was either purified by a phenol/chloroform extraction and ethanol precipitation or by the crush/soak method as described.

### Crush/soak method of RNA purification

*In vitro* transcribed RNA was run on a Novex® 6% polyacrylamide TBE-urea gel at 180 V for 30 – 45 min. The gel was removed from the plates, wrapped in plastic wrap, and placed on a TLC plate (Invitrogen). The RNA was visualized by UV shadowing with a hand-held UV lamp. The bands were excised, cut into small pieces, and two volumes of crush-soak solution were added (40) and the tubes were rotated end over end at 4°C overnight or for 2 h at room temperature. The tubes were centrifuged briefly, and the supernatant was transferred. The RNA was ethanol precipitated, washed, and resuspended in DEPC-H_2_O. RNA was quantified using the ND-1000.

## In-line probing assays

In-line probing analysis was carried out as described previously (40). The tubes were centrifuged briefly, and the supernatant was transferred. The RNA was ethanol precipitated, washed, and resuspended DEPC-H_2_O. RNA was quantified using the ND-1000.

The RNA was dephosphorylated using Calf-Intestinal Alkaline Phosphataste (NEB) and gel purified as described above. The RNA was radiolabeled with T4 PNK as described previously but with 4 µl of [gamma- ^32^P]-ATP (6000 Ci/mmol) (Perkin Elmer). The RNA was mixed with in-line buffer (40) and various amounts of metabolites as indicated. The reactions were incubated for 40 h at 25°C and then halted by the addition of 10 µl of 2X urea loading buffer. The T1 RNase ladder and alkaline hydrolysis ladder were prepared as previously described (40). The reactions were resolved by polyacrylamide gel electrophoresis using an 8% polyacrylamide/1X TBE-urea gel. The gel was dried using a gel dryer under vacuum pressure at 80°C for 90 min (FisherBiotech). The dried gel was exposed to a phosphor screen (Kodak) for 1 – 3 days and analyzed using the Typhoon™ FLA9500 Bimolecular Imager (GE Healthcare). Analysis was done using ImageQuant (GE Healthcare).

### Single-round *in vitro* transcription termination assay

Termination assays were carried out as previously described (63). The template DNA was PCR amplified from *Dv*H gDNA using standard PCR parameters. The forward primer contained the T7A1 promoter that is recognized by the *E. coli* RNA polymerase (EpiBio) and a cytosine-less leader region (DseA T7A1 prom F/R; Table S2).

Transcription was carried out in various concentrations of SAM or methionine as indicated. Products were resolved by denaturing polyacrylamide electrophoresis and visualized using the Typhoon™ FLA9500 Bimolecular Imager (GE Healthcare). Analysis was done using ImageQuant (GE Healthcare). Percent termination was determined by the amount of termination product divided by the sum of total transcription products.

### Construction of *lacZ* fusions

The predicted DseA promoter and riboswitch region were PCR amplified from *Dv*H gDNA using a forward primer with an EcoRI cut site (DseA prom/EcoRI F; Table S2) added at the 5’ end and a reverse primer with a BamHI cut site added at the 5’ end (DseA prom/ BamHI R; Table S2). The PCR product was gel purified and digested with EcoRI and BamHI for 5 min at 37°C per manufacturer’s guidelines. 1 µg of pRS415 containing a promoterless *lacZ* gene was digested with BamHI and EcoRI per manufacturer’s guidelines. The digested products were run on an agarose gel and purified as described above. The digested vector was mixed in a 1:1 molar ratio with the digested promoter and riboswitch product, 2 µl of 10X Buffer, 1 µl of T4 DNA ligase (Promega), and H_2_O up to 20 µl. Three additional reactions were carried out including a 3:1 molar ratio of vector to riboswitch, a vector only negative control, and an insert only negative control. Reactions were incubated overnight at room temperature. Tubes were placed at 65°C to halt the reaction. The plasmids were transformed into *E. coli* TOP10 cells and plated on LB plates containing ampicillin (50 µg/µl). Successful ligation and cloning was verified by PCR screening and sequencing. The new vector was named pRS415-DseA.

### β-galactosidase assays

Cells containing *lacZ* fusions were grown overnight in 5 ml of M9 minimal media with added leucine (30 mg/ml) and appropriate antibiotics. The next day, cultures were diluted 1:100 in fresh M9 minimal media in a 96-well plate and grown to an OD_600 nm_ of 0.100 – 0.200. 1 mM of methionine (Sigma) was added to half of the cultures while an equal volume of diH_2_O was added to control cultures. After 3 hours of incubation at 37°C with shaking (200 rpm) a final OD_600 nm_ was taken using a microplate reader (BioTek Synergy HT). Samples were collected for β-galactosidase measurements and were assayed as described in (64).

### Electrophoretic Gel Shift Assay (EMSA)

RNA was *in vitro* transcribed as described above. Primers were designed to amplify both the predicted sRNA DseA and a 5′ portion of the SahR (DVU0606) mRNA (DseA T7 prom F/R and SahR T7 prom F/R, Table S2). The RNA was purified by polyacrylamide gel electrophoresis and the crush-soak method. The *sahR* RNA was radiolabeled with [gamma- ^32^P]-ATP at the 5′ end after dephosphorylation by CIP. 0.4 pmol of DseA RNA or control RNA (Dv SIC2, generated using primers Dv sRNA-2 T7 prom F/R) was mixed with 0.2 pmol of end-labeled SahR RNA in 5 µl of binding buffer (65). The mixture was incubated at 70°C for 5 min and then at 37°C for 20 min. Loading buffer II (Ambion) was added and the samples were loaded onto a Novex^®^ 6% TBE gel. The gel was run at 200 V for 30 – 45 min. The gel was removed from the plates and vacuum dried on Whatman™ paper. The gel was exposed to Fuji film overnight at −80°C. The film was developed manually by brief immersion in Kodak developer and fixer.

### Construction of a DseA deletion and complement strain

The deletion strain, Δ*dseA*, was constructed by the J. Wall Laboratory (University of Missouri) as described in Bender et al. (66,67). A region upstream of DseA, the neomycin phosphotransferase (*npt*) gene that confers resistance to kanamycin and G418, and a region downstream of DseA were PCR amplified and then fused together via overhangs into one product similar to previously described protocols. This product was ligated into an *E. coli* cloning vector, which is not stable in *Dv*H, and transformed into *E. coli* TOP10 cells. This vector was then electroporated into *Dv*H as described previously (67). Transformants were screened and sequenced to verify the deletion of DseA by homologous recombination and the new strain was designated Δ*dseA*.

To complement the Δ*dseA* strain, the region corresponding to the DseA +1 site through the terminator region (*Dv*H coordinates 1,264,319 – 1,264,156) was amplified with primers DseA-pSIL300-BamHI-F/DseA-pSIL300R. This allowed for directional cloning into the pSIL300 vector, which is a derivative of pMO719 (68) that possesses the promoter for the *Dv*H cytochrome *c*_3_ gene (DVU3171) with BamHI and ScaI sites (GAGTCCCAAACCGCCATGAATCTAGGCTTTCCCGCTCCATTCCTTGACACTCTATCATTGATAGAGTTACCATCCCGCTCCCTATCAGTGATAGAGAGG<u>GGGATCC</u>ATAT<u>AGTACT</u>AATA). This cytochrome *c*_3_ promoter was inserted into the EcoRV site of the parent vector using primers Xba-c3pro-F and c3proBamSca-R. The resulting plasmid, p*dseA*, was transformed into the Δ*dseA* strain as described above and the complement strain was selected by plating on LS4D containing both G418 and spectinomycin.

## Acknowledgements

We thank Dr. Judy Wall and Tom Juba (University of Missouri) for construction of the deletion strain. We thank Dr. Derek Fisher (Southern Illinois University Carbondale) for assistance with the β-galactosidase assays and helpful discussions.

---

**Figure S1.** Alignment of RACE clone sequences corresponding to DseA. Positions 1-25 correspond to the 3’-end and position 26 corresponds to the +1 site of the RNA as depicted in Figure 1B.

**Figure S2.** Alignment of conserved DseA sequences. Abbreviations are as follows: DvH, *Desulfovibrio vulgaris* Hildenborough; Dv RCH1, *Desulfovibrio vulgaris* RCH1; Dv DP4, *Desulfovibrio vulgaris* DP4; DvM, *Desulfovibrio vulgaris* Miyazaki F; Ds 2638, *Desulfovibrio salexigens* DSM2638; Da G20, *Desulfovibrio alaskensis* G20. Black shading indicates identically conserved bases while grey shading represents similarly conserved bases.

**Figure S3.** Alignment of the predicted riboswitch region of two *Desulfovibrio* species and other known SAM riboswitches. Alignment was generated by Rfam (48) and the colors represent the consensus base for that location.

**Figure S4.** RT-PCR analysis of DVU1170 region. (**A**) Co-transcription of DseA and DVU1170. Top of panel illustrates the genomic view of the DseA-DVU1170 locus with lines a, b, c, and d indicating regions of the locus targeted by RT-PCR. Primers 1170 RT F1-F4/R (Table S2) were used to target regions a-d. (**B**) DVU1170 and DVU1169 are not co-transcribed. Top of panel illustrates the genomic view of the DVU1170-DVU1169 locus with lines a, b, and c indicating regions of the locus targeted by RT-PCR. Primers DVU1169 RT F1-3/R were used to target regions a-c. Gel analysis of the RT-PCR results are provided in the bottom of each panel. The reactions within each set of four wells corresponding to the mapped genome regions are as follows: (−), PCR without DNA template as a negative control; (−RT), PCR with RNA as the template as a negative control; (+), PCR with genomic DNA from *Dv*H as a control; and (+RT), RT-PCR with RNA as a template.

**Figure S5.** DseA-*sahR* mRNA interaction region. (**A**) The sequence encompassing the −36 to +126 (in reference to the start codon) region of the *sahR* mRNA. The RBS is underlined, and the start codon is bolded. The predicted region of interaction between DseA and *sahR* is shown in red. (**B**) EMSA showing interaction between DseA and RNA oligos of portions of the *sahR* mRNA. The sequence of oligo 1, oligo 2, and oligo 3 is underlined with a black, blue, and purple line, respectively. The aforementioned oligos were generated using the following primer sets in Table S2: SahR T7 prom F/SahR Rev 2 (oligo 1), SahR T7 prom F/SahR Rev 3 (oligo 2), and SahR Middle F/SahR Rev 3 (oligo3).

