## Supplemental Materials for "A dual functioning small RNA/Riboswitch controls the expression of the methionine biosynthesis regulator SahR in *Desulfovibrio vulgaris* Hildenborough"

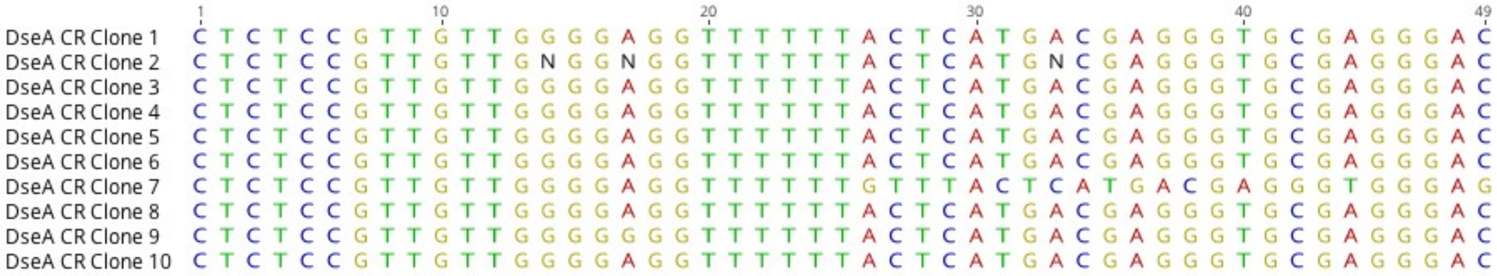


**Figure S1.** Alignment of RACE clone sequences corresponding to DseA. Positions 1-25 correspond to the 3'-end and position 26 corresponds to the +1 site of the RNA as depicted in Figure 1B.

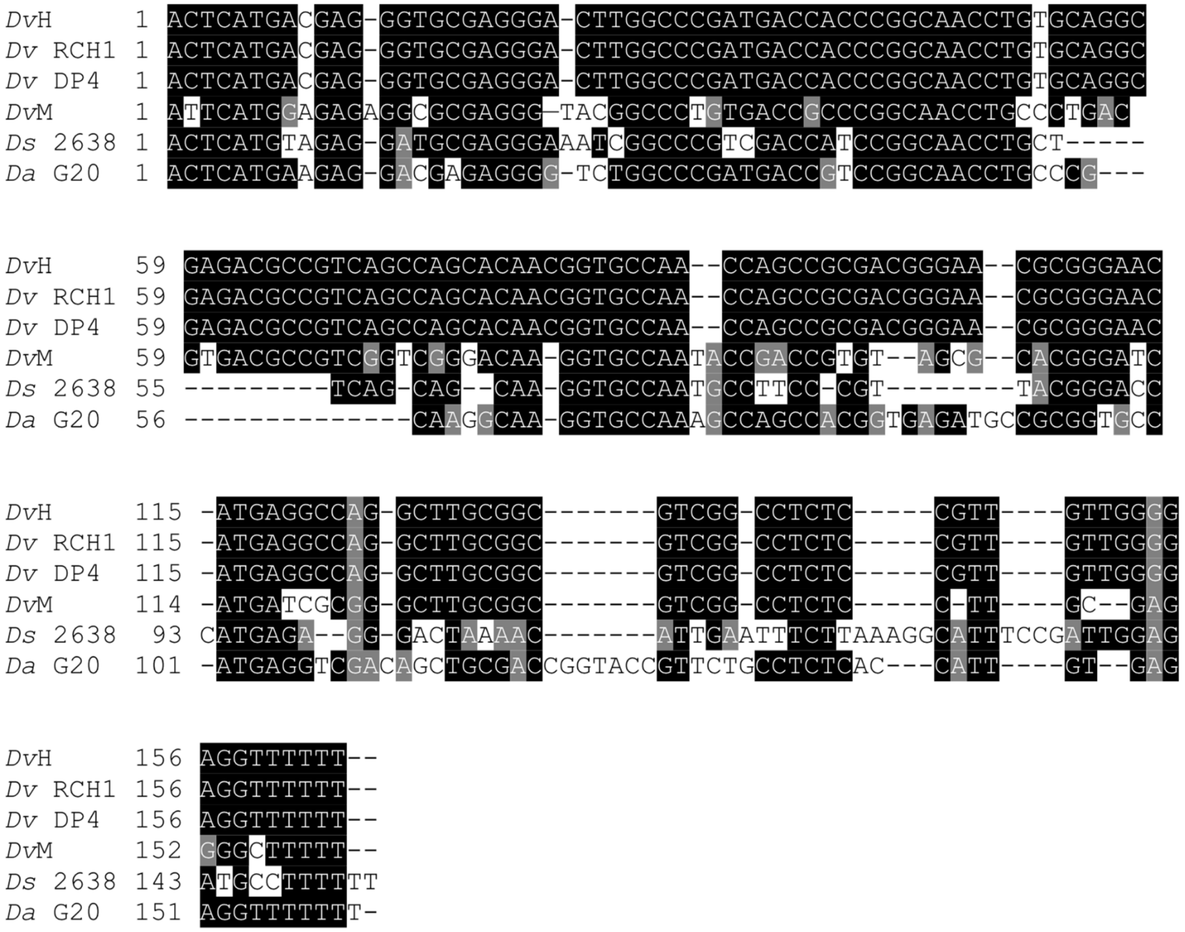


**Figure S2.** Alignment of conserved DseA sequences. Abbreviations are as follows: DvH, *Desulfovibrio vulgaris* Hildenborough; Dv RCH1, *Desulfovibrio vulgaris* RCH1; Dv DP4, *Desulfovibrio vulgaris* DP4; DvM, *Desulfovibrio vulgaris* Miyazaki F; Ds 2638, *Desulfovibrio salexigens* DSM2638; Da G20, *Desulfovibrio alaskensis* G20. Black shading indicates identically conserved bases while grey shading represents similarly conserved bases.

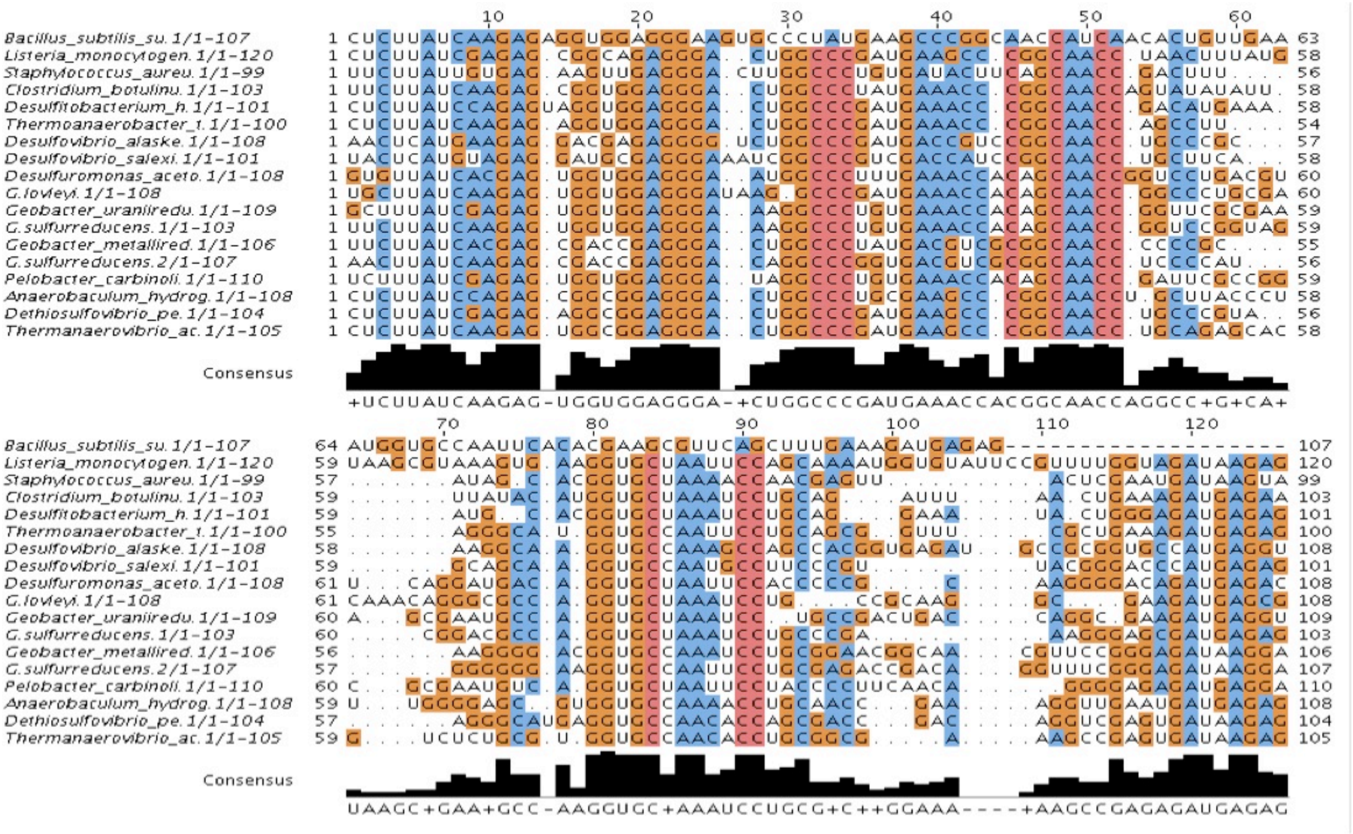


**Figure S3.** Alignment of the predicted riboswitch region of two *Desulfovibrio* species and other known SAM riboswitches. The colors represent the consensus base for that location.

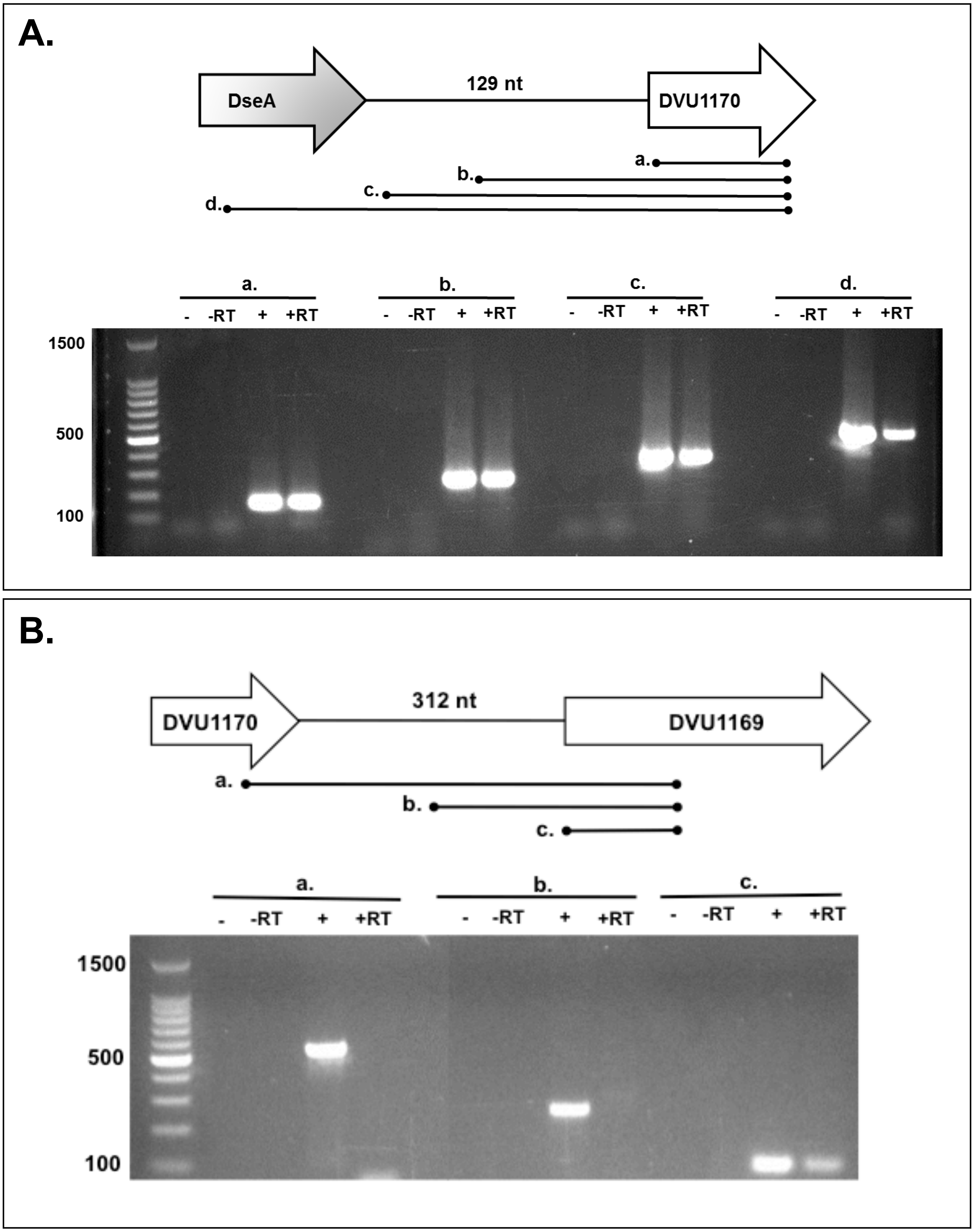


**Figure S4.** RT-PCR of DVU1169 transcript. **(A)** Genomic view of the DVU1170 and DVU1169 region. Lines a, b, and c indicate regions of the locus amplified by PCR as shown in panel B. **(B)** RT-PCR results. Lanes 1–4 corresponds to line a, lanes 5–8 to line b, lanes 9–12 to line c. Within each set of 4 wells, PCR to amplify lines a–c was performed with RNA after RT (+RT), RNA without RT as a control (-RT), PCR without DNA template as a control (-), and PCR with genomic DNA from *Dv*H as a positive control (+).

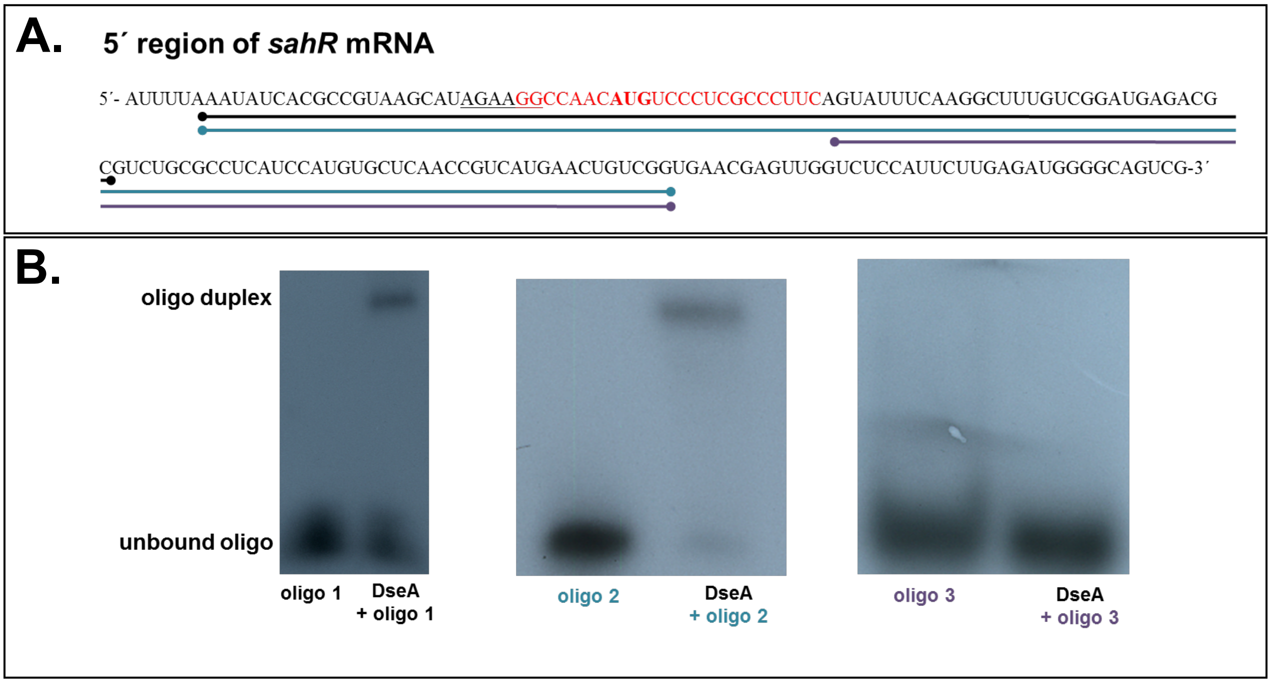


**Figure S5. (A)** The sequence encompassing the -36 to +126 (in reference to the start codon) region of the *sahR* mRNA. The RBS is underlined and the start codon is bolded. The predicted region of interaction between DseA and sahR is shown in red. **(B)** EMSA showing interaction between DseA and RNA oligos of portions of the *sahR* mRNA. The sequence of oligo 1, oligo 2, and oligo 3 is underlined with a black, blue, and purple line, respectively.

**TABLE S1.** Bacterial strains and plasmids used in this study.

| **Strains** | **Genotype/Description** | **Source/Reference** |
|---|---|---|
| *Desulfovibrio vulgaris* Hildenborough | Wild-type strain Hildenborough | ATCC 29579$^T$ |
| Δ*dseA* | *Dv*H, Δ*dseA*, Km$^R$, G418$^R$ | Judy Wall Lab (University of Missouri) |
| *E. coli* TOP10 | F- *mcrA* Δ( *mrr-hsd*RMS-*mcr*BC) Φ80*lac*ZΔM15 Δ *lac*X74 *rec*A1 *ara*D139 Δ( *araleu*)7697*gal*U *gal*K *rps*L (StrR) *end*A1 *nup*G | Invitrogen |
| **Plasmids** | | |
| pRS415 | *E. coli* expression vector, promoterless *lacZ*, Amp$^R$ | Simons et al., 1983 |
| pRS415-DseA | pRS415 with the promoter region of DseA, Amp$^R$ | This study |
| p*dseA* | pSIL300 with DseA, Spec$^R$ | This study |

**TABLE S2.** Primers and probes used in this study.

| Name | Sequence (5′ – 3′) |
|---|---|
| DseA 30mer Probe | GCAAGCCTGGCCTCATGTTCCCGCGTTCCCGTCGC |
| DvH 5S 40mer Probe | TTTCCCACGAGCTACCCCGCAGTATCATCGGCGATGGAGGGC |
| DseA CR F | AACATGAGGCCAGGCTTGCGG |
| DseA CR R | GTTCCCGTCGCGGCTGGTTG |
| 1170 RT F1 | ATGACCACCCGGCAACCTGTGCAG |
| 1170 RT F2 | CACATGTCGTGGCTGCAGGTGGGT |
| 1170 RT F3 | GCCACGCGCATAGGGGAACACGATG |
| 1170 RT F4 | ATGCCCGTCCAATGGACCGATCCC |
| 1170 RT R | TGTTCTTGCGGAAGGCTCGGCCATT |
| DVU1169 RT F1 | AAGGAGCGGAGACGGTCTGGAATGG |
| DVU1169 RT F2 | TCTGCACCGCATCAACGCGACTCC |
| DVU1169 RT F3 | TTCTCCGTCATCTGCGTCCTGTCTCTCG |
| DVU1169 RT R | GCCTCGCCGAACGATTCGTTGTAGTCC |
| DseA prom/EcoRI F | GAATTCGTTGACGTTTTTTCTGCGACC |
| DseA prom/ BamHI R | GGATCCACGACATGTGTCGTGCTG |
| 16S qRT-PCR F2 | GTGCGAAAGCGTGGGGAGCA |
| 16S qRT-PCR R2 | ACGGCACCGAAGCTCAAGGC |
| DVU1170 qRT-PCR F | AGATGCGACGCAGATGGACGAATTGC |
| DVU1170 qRT-PCR R | ATGATGTTCTTGCGGAAGGCTCGGC |
| sahR qRT-PCR F | GCCGATTTCGCCAGCATCAACATGG |
| sahR qRT-PCR R | CGAAATCCGTGACCATGAGTTGGCCT |
| DseA T7 prom F | TAATACGACTCACTATAGGGACTCATGACGAGGGTGCGAGGGAC |
| DseA T7 prom R | CAAAAAACCTCCCCAACAACGGAGAGGCC |
| DseA R | GGCATGGGCGCTTCCTCGTTGTGGTA |
| DseA T7A1 prom F | TTGACTTAAAGTCTAACCTATAGGATACTATGTAGTAAGGAGGTTGTAT GGAAGAACTCATGACGAGGGTGCGA |
| DseA T7A1 R | CGTGCAACGGCCTGAAATCCACCCACCT |
| SahR T7 prom F | TAATACGACTCACTATAGGGTAAATATCACGCCGTAAGCATAGAA |
| SahR T7 prom R | GCAAACCGGCACCCGTGAGAATCTT |
| SahR Middle F | TAATACGACTCACTATAGGGAGTATTTCAAGGCTTTGTCGGAT |
| SahR Rev 2 | GCGTCTCATCCGACAAAGCCTTGAAATACT |
| SahR Rev 3 | CCGACAGTTCATGACGGTTGAGCACATG |
| Dv sRNA-2 T7 prom F | TAATACGACTCACTATAGGGAGGAGCCGGTTCTTGATGGTCAT |
| Dv sRNA-2 T7 prom R | CCCGGAGGCTTCGCAGGAAGGAACA |
| DVU0277 qRT-PCR F | GACACCCTGCATCGTTATGA |
| DVU0277 qRT-PCR R | CCCATCACAGTGGAGAAGATAC |
| DVU1165 qRT-PCR F | GAAGGATGTGGAGGTGATGTT |
| DVU1165 qRT-PCR R | CTATCTGTTCCGGCTCAAGTT |
| DVU1362 qRT-PCR F | GCTTCTCTCCCGTGTTCTC |

| | |
|---|---|
| DVU1362 qRT-PCR R | GCCAGTATCTTGACCGTGAT |
| DVU1411 qRT-PCR F | TCCGTCCCACCCTTCTT |
| DVU1411 qRT-PCR R | CACCTTCTGCATTTCGCATTC |
| DVU1977 qRT-PCR F | GGCGACATGGTGCTGTT |
| DVU1977 qRT-PCR R | TACTCGATGATGGCGAGGAT |
| DVU 2373 qRT-PCR F | CATCTGGGATCTCGGCTATTTC |
| DVU2373 qRT-PCR R | GAGCCTTCGATGACGATGTT |
| DVU2514 qRT-PCR F | CATCTTCCGCCTCAACTTCT |
| DVU2514 qRT-PCR R | CGATTTCGCCTATCCGTATCTT |
| DVU2938 qRT-PCR F | CCCACCATCAAGACGTCGAA |
| DVU2938 qRT-PCR R | TCCGGGGTAGACCTTGTTCA |
| DVU3156 qRT-PCR F | AAATCCAACGACGGTTCTCTC |
| DVU3156 qRT-PCR R | TCTCGGAGTGGAAGACGTAG |
| rplS qRT-PCR F from ref. (59) | TGTCTTCCCCCTGCACTCG |
| rplS qRT-PCR R from ref. (59) | CTTGATGCGGGCAGCCTTAC |
| Xba-c3pro-F | GAGCTCTAGAGTCCCAAACCGCCATGAATCTAG |
| c3proBamSca-R | TATTAGTACTATATGGATCCCCCAAGCGGGATGGTATTGTGTC |
| DseA-pSIL300-BamHI-F | AATAGGATCCACTCATGACGAGGGTGCGAG |
| DseA-pSIL300-R | AAAAAACCTCCCCAACAACG |

**TABLE S3.** IntraRNA predicted targets of DseA.

| Target | Energy kcal/mol | Gene | Annotation |
|---|---|---|---|
| DVU2373 | -25.6163 | | OMP85 family outer membrane protein |
| DVU0277 | -24.8905 | | AraC family transcriptional regulator |
| DVU1165 | -21.5161 | | pyridine nucleotide-disulfide oxidoreductase |
| DVU2594 | -21.4385 | | |
| DVU1977 | -19.828 | *groES* | co-chaperonin GroES |
| DVU1362 | -19.4161 | | |
| DVU3156 | -19.2118 | *glmS* | glucosamine--fructose-6-phosphate aminotransferase (isomerizing) |
| DVU0606 | -19.1926 | | ArsR family transcriptional regulator |
| DVU2514 | -19.1684 | *pyk* | pyruvate kinase |
| DVU2605 | -18.978 | | |
| DVU1411 | -18.8335 | *thiC* | thiamine biosynthesis protein ThiC |
| DVU1902 | -18.7894 | | |
| DVU2623 | -18.7117 | | |
| DVU2620 | -18.6032 | | |
| DVU0297 | -17.9608 | | |
| DVU1923 | -17.919 | *hupD* | hydrogenase expression/formation protein HupD |
| DVU2282 | -17.8726 | | |
| DVU1824 | -17.7374 | | |
| DVU0917 | -17.5822 | *atpE* | ATP synthase F0 subunit C |
| DVU2331 | -17.4978 | | Smr family protein |
| DVU2938 | -15.539 | | DUF39, predicted homocysteine formation from aspartate semialdehyde |
